# Age and sex dependent effects of metabolic response to muscle contraction

**DOI:** 10.1101/2023.05.30.542769

**Authors:** Matthew D. Campbell, Danijel Djukovic, Daniel Raftery, David J. Marcinek

**Affiliations:** Department of Radiology, University of Washington, Seattle, WA; Anesthesiology & Pain Medicine, University of Washington, Seattle, WA

**Keywords:** high-intensity intervals, low-intensity steady state, sarcopenia, metabolism, age, sex

## Abstract

Sarcopenia, the age-related loss of muscle mass and function, contributes to decreased quality of life in the elderly and increased healthcare costs. Decreased skeletal muscle mass, specific force, increased overall fatty depositions in the skeletal muscle, frailty and depressed energy maintenance are all associated with increased oxidative stress and the decline in mitochondrial function with age. We hypothesized that elevated mitochondrial stress with age alters the capacity of mitochondria to utilize different substrates following muscle contraction. To test this hypothesis, we designed two *in vivo* muscle-stimulation protocols to simulate high-intensity intervals (HII) or low intensity steady-state (LISS) exercise to characterize the effect of age and sex on mitochondrial substrate utilization in skeletal muscle following muscle contraction. Following HII stimulation, mitochondria from young skeletal muscle increased fatty acid oxidation compared to non-stimulated control muscle; however, mitochondria from aged muscle decreased fatty acid oxidation. In contrast, following LISS, mitochondrial from young skeletal muscle decreased fatty acid oxidation, whereas aged mitochondria increased fatty acid oxidation. We also found that HII can inhibit mitochondrial oxidation of glutamate in both stimulated and non-stimulated aged muscle, suggesting HII initiates circulation of an exerkine capable of altering whole-body metabolism. Analyses of the muscle metabolome indicates that changes in metabolic pathways induced by HII and LISS contractions in young muscle are absent in aged muscle. Treatment with elamipretide, a mitochondrially targeted peptide, restored glutamate oxidation and metabolic pathway changes following HII suggesting rescuing redox status and improving mitochondrial function in aged muscle enhances the metabolic response to muscle contraction.

**Statements and Declarations:** The authors declare no competing financial interests.

## Introduction

Metabolic response to exercise is a complicated process that involves multiple energy systems [1]. The basic energetic currency of cells is ATP and through various pathways and mechanisms ATP is synthesized to meet ATP demand during exercise and recovery [2]. During short bouts of intense exercise PCr and glycogen are both mobilized to address energetic demands [3, 4]. With longer bouts of exercise and during recovery following muscle contraction ATP generation by mitochondrial oxidative phosphorylation is central to maintaining and restoring energy homeostasis. Aging skeletal muscle is associated with decreased mitochondrial ATP production and elevated oxidant production leading to disruptions in both humans and rodents [5–9], although there continues to be some debate about whether the decline in the capacity for mitochondrial ATP production in human skeletal muscle is an intrinsic part of aging or is driven by secondary factors like reduced physical activity [10]. However, when mitochondrial respiratory capacity is analyzed in the context of mitochondrial content or sub-maximal metabolism evidence supports a decline in mitochondrial quality in aging skeletal muscle that contributes to disruption of both energy and redox homeostasis under resting conditions [5, 6, 9, 11]. It is also still not clear how this elevated mitochondrial induced stress impairs mitochondrial ability to respond to increased energetic demand of muscle contraction[12, 13].

Metabolomics can be used as a powerful complement to direct measurement of mitochondrial function. Recent work in metabolomics has made great strides in examining both response to exercise [14] and aging [15, 16]. However, one drawback with many metabolomic studies of exercise is that it is performed on circulating metabolites. This necessarily omits crucial information about metabolism within the tissue itself. Although muscle metabolism is partially dependent on circulating metabolites [2] many of the mechanisms of mitochondrial substrate utilization during and following exercise are still misunderstood. Importantly there is relatively little known about how aging effects metabolomic response to exercise in skeletal muscle. This study was designed as a multilevel approach using metabolomics and specific tests of mitochondrial substrate utilization to test both sex and age dependent effects of the metabolic response to muscle contraction. We designed this study to test the hypothesis that age inhibits mitochondrial substrate utilization following exercise.

## Results

### Sex and age differences in response to high-intensity stimuli

To test potential sex differences in response to muscle contraction with age we designed two *in vivo* muscle contraction protocols. Young and aged male and female mice were stimulated with either low-intensity steady-state (LISS) or high-intensity interval (HII) contraction protocols. Aged female mice generate lower total force as measured by area under the curve during HII compared to young female mice (Figure 1A). However, male mice showed no difference during HII (Figure 1B). There were no differences with age in both females and male mice during LISS contraction (Figure 1 C & D). Both female and male aged mice had lower peak force during HII contraction and aged female mice had lower force compared to aged males (Figure 2A). There was no effect of age or sex on max force during LISS contractions (Figure 2B).

**Figure 1.**
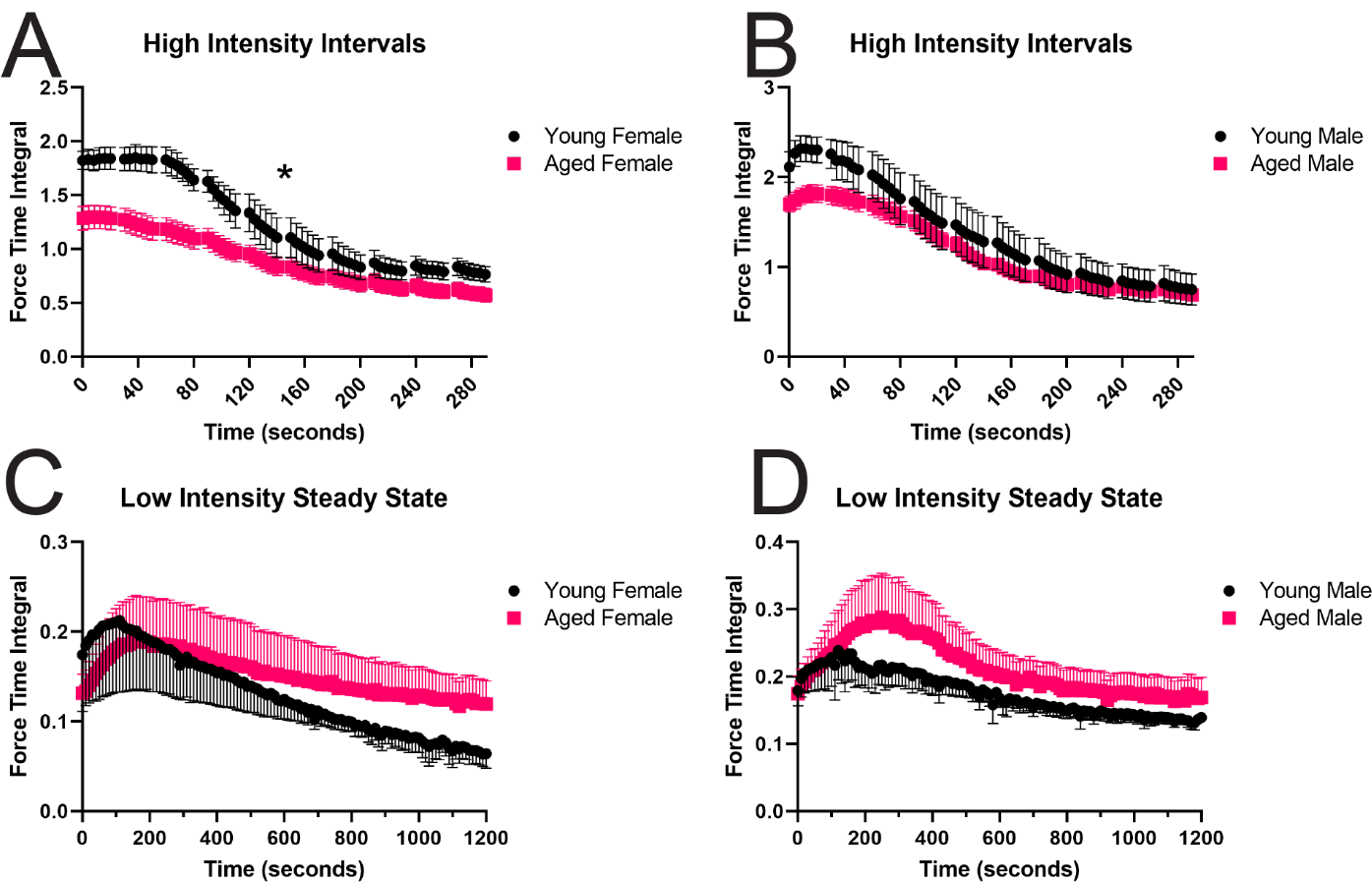
Force generation during contraction protocols. **A.** High-intensity intervals in female mice, n=5-7 **B.** High-intensity intervals in male mice, n=5-9 **C.** Low-intensity steady state in female mice, n=5-8 **D.** Low-intensity steady state in male mice, n=5-8. All data represented as mean ± S.E.M., *-p<0.05, two-way ANOVA effect of age.

**Figure 2.**
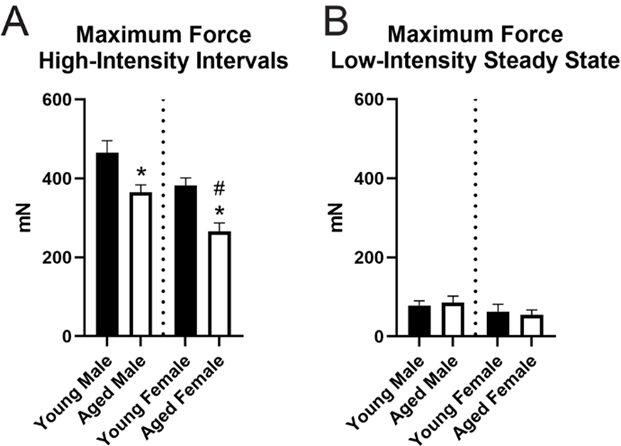
Maximum force generation during contraction protocols. **A.** High-intensity intervals, n=5-9 **B.** Low-intensity steady state, n=5-8. *-p<0.05 One-way ANOVA effect of age, all data represented as mean ± S.E.M., #-p<0.05 one-way ANOVA effect of sex

### Sex and age differences in substrate utilization following muscle contraction

To test mitochondrial substrate utilization, we measured respirometry in permeabilized red gastrocnemius following contraction. We used both the contracted (stimulated) leg and the non-stimulated leg for an internal control comparison. Additionally, we sacrificed non-stimulated naïve animals to test for systemic response to acute muscle contraction. Following HII, young female gastrocnemius did not have significantly different glutamate utilization between contracted and non-stimulated muscle, however both contracted and non-stimulated muscle utilized significantly more glutamate than naïve muscle (Figure 3A). Following HII, aged female gastrocnemius did not oxidize glutamate in either the contracted or non-stimulated muscle and both were significantly decreased compared to naïve muscle (Figure 3B). Similar to the HII, LISS significantly increased glutamate utilization in both the contracted and non-stimulated muscle of young female muscle compared to naïve (Figure 3C). Unlike HII, LISS increased glutamate utilization in aged female muscle compared to the non-stimulated and both contracted and non-stimulated muscle had significantly greater glutamate oxidation than naïve muscle (Figure 3D).

**Figure 3.**
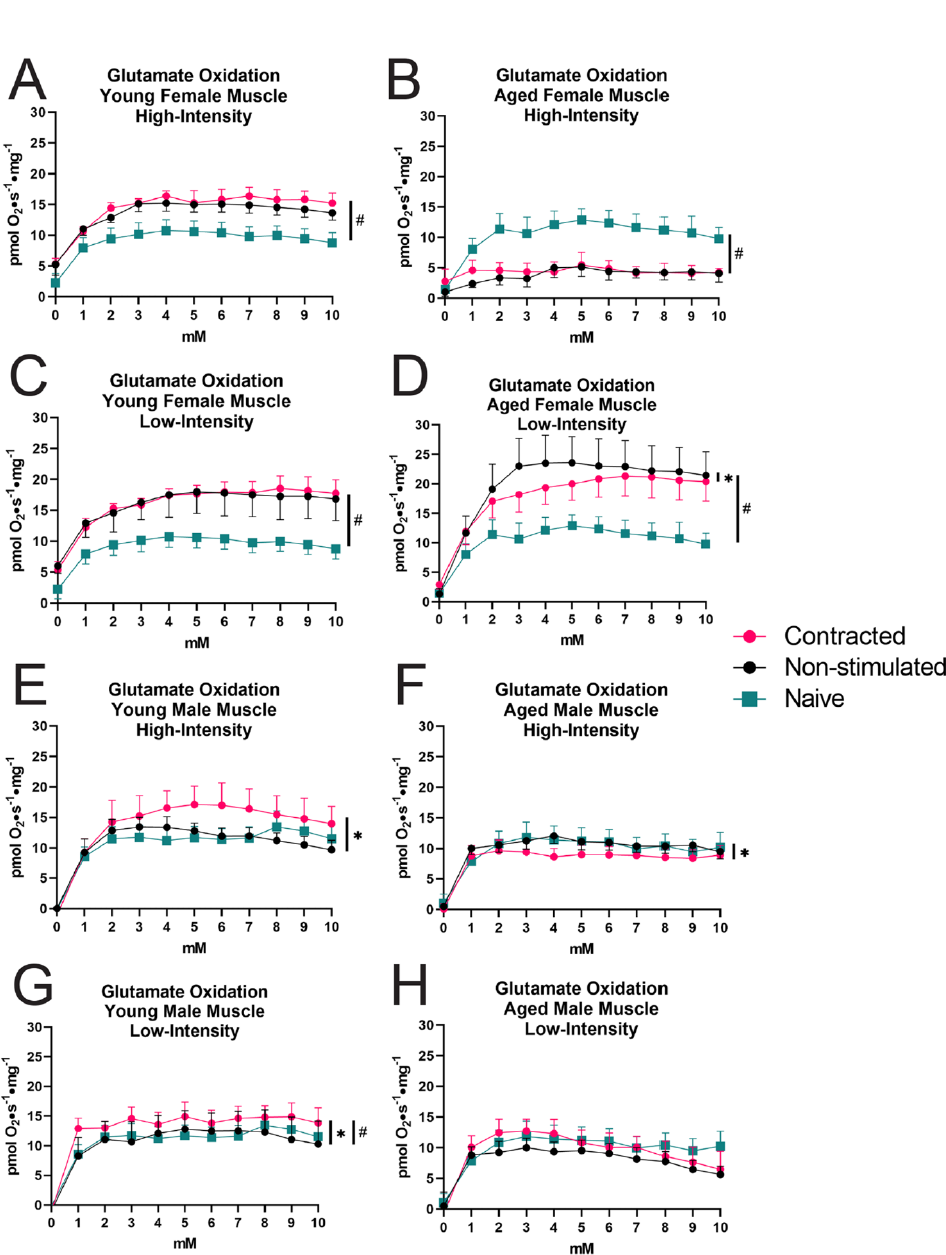
Glutamate utilization following acute muscle contraction. **A.** High-intensity intervals in young female mice, n=5 **B.** High-intensity intervals in aged female mice, n=7 **C.** Low-intensity steady state in young female mice, n=5 **D.** Low-intensity steady state in aged female mice, n=8 **E.** High-intensity intervals in young male mice, n=5 **F.** High-intensity intervals in aged male mice, n=9 **G.** Low-intensity steady state in young male mice, n=5 **H.** Low-intensity steady state in aged male mice, n=8. All data represented as mean ± S.E.M., *-p<0.05 repeated measures one-way ANOVA stimulated compared to non-stimulated, #-p<0.05 repeated measures one-way ANOVA effect compared to naïve.

The response to stimulation was sex-specific with more subtle effects in the males than observed in the females. HII stimulation increased glutamate oxidation in the stimulated leg only in young male mice (Figure 3E), but slightly decreased glutamate oxidation in aged male mice (Figure 3F). However, LISS increased glutamate oxidation in young male mice (Figure 3G) with no change in aged males (Figure 3H). Fat metabolism has been shown to be different between females and males in both rodents [17] and humans [18] so we also tested fatty acid utilization following high and low intensity contraction. Compared to glutamate oxidation, the skeletal muscle mitochondria demonstrated a low capacity to oxidize fatty acids. HII increased fatty acid oxidation compared to non-stimulated legs in young females (Supplemental Figure 1A) but decreased respiration in aged female mice (Supplemental Figure 1B). Whereas LISS decreased respiration compared to non-stimulated legs in young females (Supplemental Figure 1C) but increased in aged females. Male muscle did not show differences in respiration between the contracted and non-stimulated muscle using fatty acids except in young male muscle following LISS (Supplemental Figure 1 E-H).

### Age decreases metabolic pathway changes following muscle contraction

Metabolite levels change following exercise in both mice [19, 20], and humans [21, 22]. To date most studies have examined the metabolite changes of serum following whole-body exercise. We were interested in the muscle-specific metabolic changes, so we performed targeted metabolomics on the gastrocnemius muscles frozen immediately after contraction to compare the contracted muscle to the non-stimulated muscle and examine changes with age. We measured between 180-186 of the 300 targeted aqueous metabolites in each sample (Supplemental Table 1). Following HII contraction we found 22 metabolite levels changed in young muscle (Figure 4A) and only eight metabolite levels changed in aged muscle (Figure 4B) using paired analysis of the stimulated versus non-stimulated muscle. Of the eight significant metabolites measured in aged muscle following HII 6 are also present as significantly altered metabolites in the young comparison: glucosamine 6-phosphate, glucose 1-phosphate, 3-hydroxybutyric acid, sorbitol, arginosuccinic acid, and alpha-hydroxyisobutyric acid. Using all measured metabolites, we analyzed pathway changes including correction for multiple testing following HII in young muscle (Figure 5A) and aged muscle (Figure 5B). Following HII, 20 pathways were significantly altered relative to young non-stimulated muscle (Figure 5C), however in aged muscle only five pathways were significantly different after contraction (Figure 5D). Following LISS contraction we found 17 metabolites significantly affected by contraction in young muscle (Figure 6A) and only seven metabolites changed in aged muscle (Figure 6B). Of the seven significant metabolites measured in aged muscle following LISS none are also present as significantly altered metabolites in the young comparison. Using all measured metabolites, we analyzed pathway changes following LISS in young muscle (Figure 7A) and aged muscle (Figure 7B). Following LISS 16 pathways were significantly changed compared to young non-stimulated muscle (Figure 7C), however in aged muscle only three pathways were significantly changed (Figure 7D).

**Figure 4.**
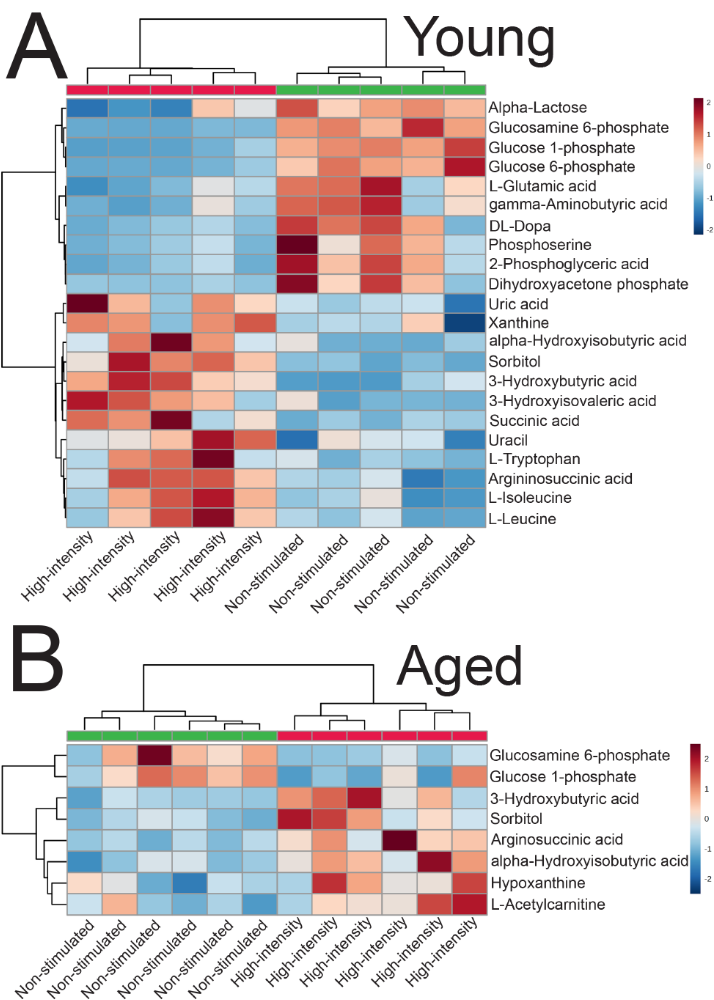
Significant metabolite changes following high-intensity contractions. **A.** Young females, n=5 **B.** Aged females, n=6. Stimulated muscle shown in red and non-stimulated in green on the dendrogram. All data analyzed by paired students t-test, p<0.05.

**Figure 5.**
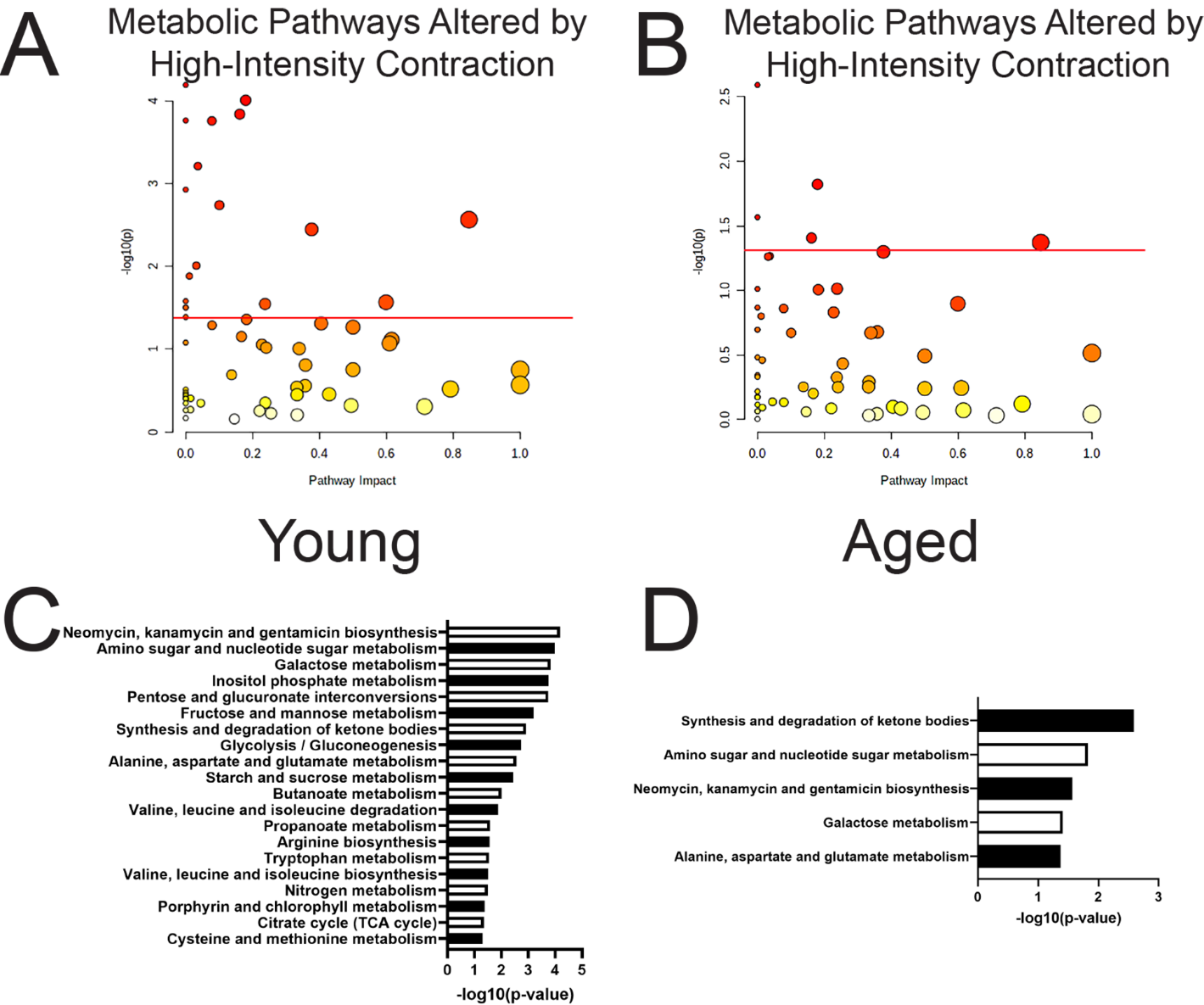
Metabolic pathways altered by high-intensity contractions. **A.** Young females, n=5, and **B.** Aged females, n=6. Pathway impact represents the number of metabolites in each pathway that are significantly altered. **C.** Individual pathways altered by high-intensity contractions and the corresponding p-value in young females, and **D.** Aged females.

**Figure 6.**
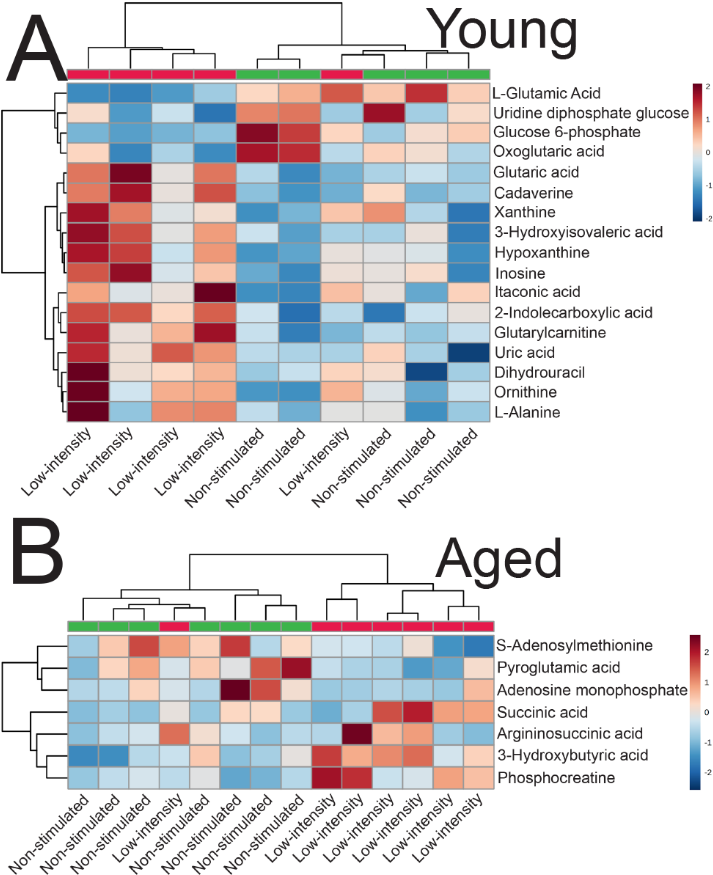
Significant metabolite changes following low-intensity contractions. **A.** Young females, n=5 **B.** Aged females, n=7. Stimulated muscle shown in red and non-stimulated in green on the dendrogram. All data analyzed by paired students t-test, p<0.05.

**Figure 7.**
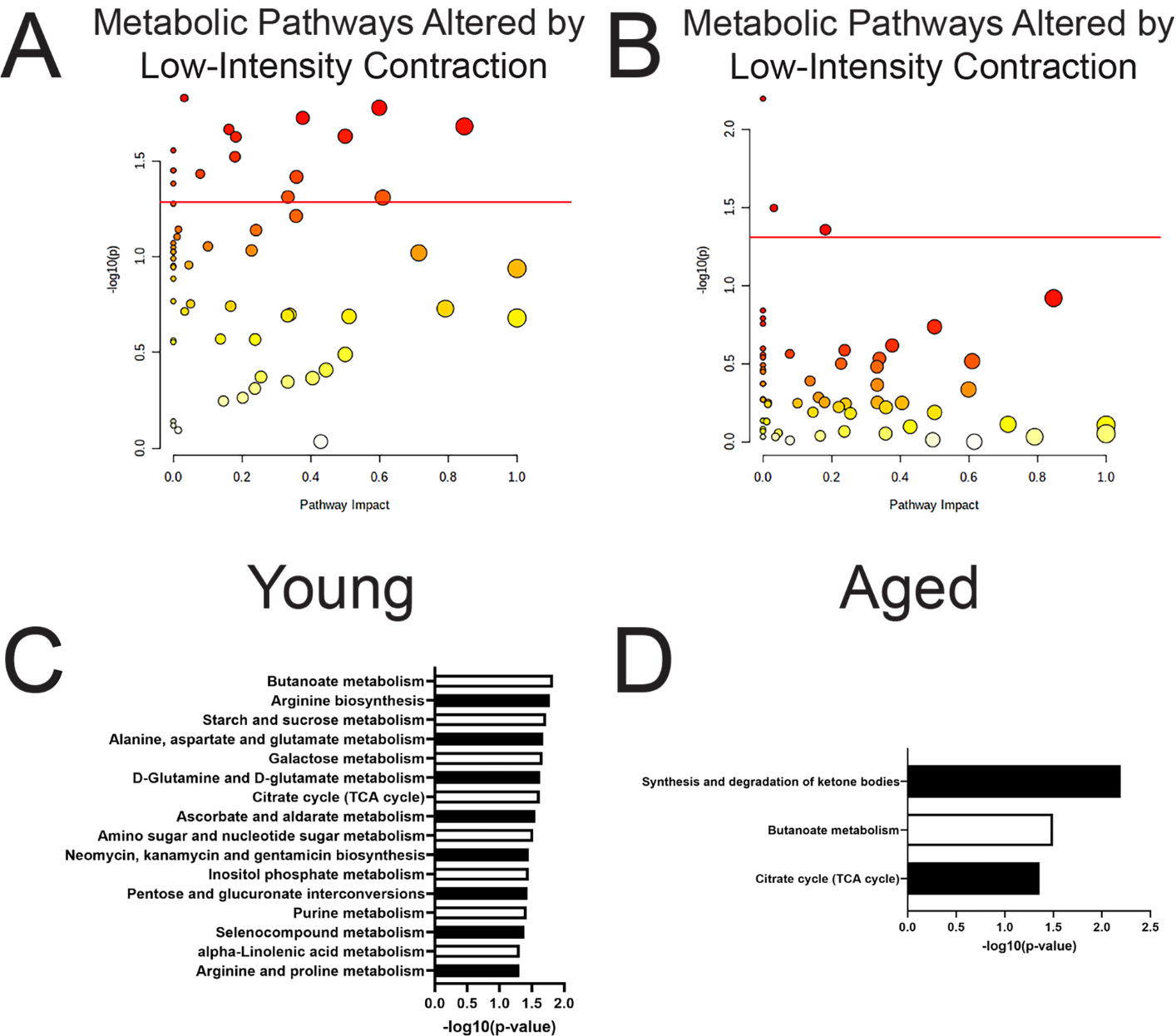
Metabolic pathways altered by low-intensity contractions. **A.** Young females, n=5. Pathway impact represents the number of metabolites in each pathway that are significantly altered. **B.** Aged females, n=7. Pathway impact represents the number of metabolites in each pathway that are significantly altered. **C.** Individual pathways altered by low-intensity contractions in young females and their corresponding p-value. **D.** Individual pathways altered by low-intensity contractions in aged females and their corresponding p-value.

### Elamipretide rescues metabolic response to high-intensity contraction in aged muscle

We have previously shown that elamipretide (ELAM, formerly known as SS-31 and Bendavia) increases fatigue resistance and improves mitochondrial bioenergetics in aged skeletal muscle [8, 9]. Further we have shown that ELAM interacts directly with proteins involved in glutamate metabolism [23]. To test if treatment with ELAM restored glutamate oxidation or the response of metabolic pathways to contraction, we treated aged females with ELAM for 8 weeks. Following treatment, we performed HII stimulation. There was no difference in overall fatiguability (Figure 8A), or force production (Figure 8B) compared to untreated aged females. ELAM restored glutamate utilization following HII in both the contracted and non-stimulated legs (Figure 8C). Additionally following HII contraction we found 24 metabolites changed in aged ELAM treated muscle compared to non-stimulated muscle (Figure 8D). We also used all measured metabolites to analyze metabolic pathway changes between contracted and non-stimulated muscle (Figure 8E) and found that 13 metabolic pathways were significantly altered by HII contraction (Figure 8F).

**Figure 8.**
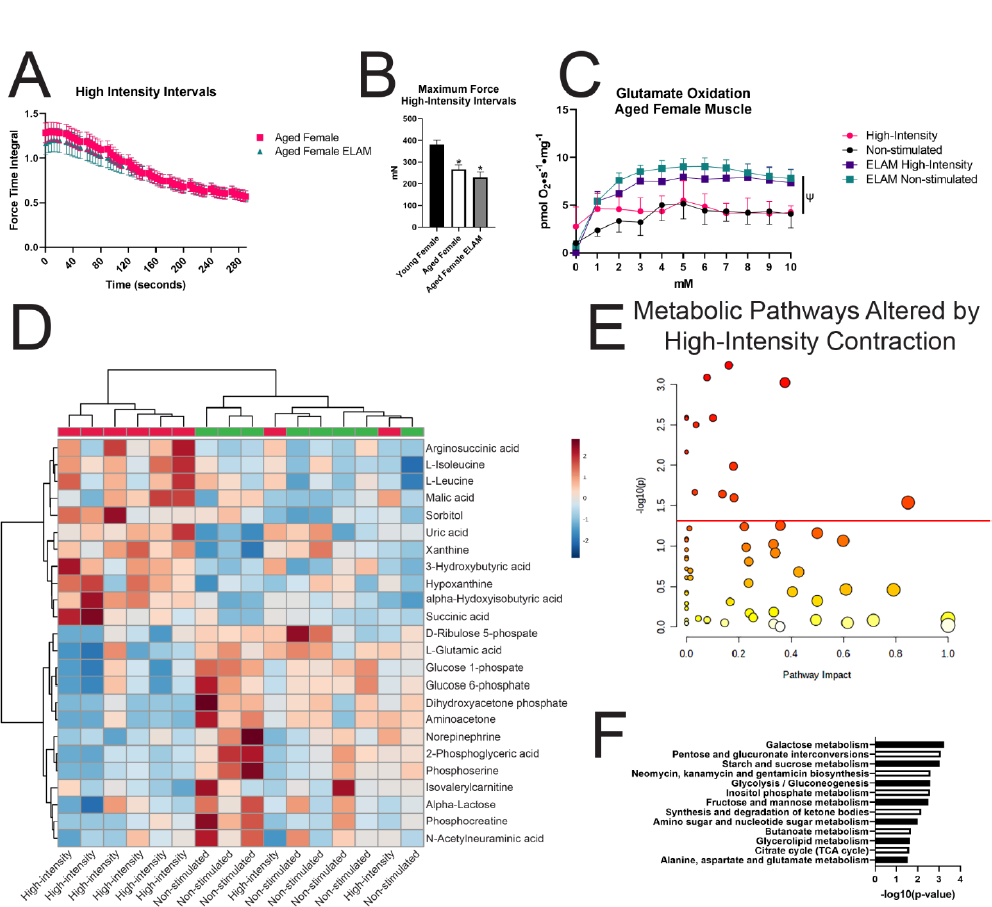
Elamipretide effects in aged female mice using high-intensity intervals. **A.** Force generation during high-intensity intervals, n=8 **B.** Maximum force generation during high-intensity intervals, n=5-8, *-p<0.05 one-way ANOVA compared to young female **C.** Glutamate utilization following high-intensity intervals, n=7-8, Ψ-p<0.05 repeated measures one-way ANOVA effect compared to aged high-intensity **D.** Significantly altered metabolites between contracted and non-stimulated gastrocnemius, n=8 **E.** Comparison of all measured metabolic pathway changes between contracted and non-stimulated gastrocnemius, n=8 **F.** All significantly altered metabolic pathways, n=8, students paired t-test.

## Discussion

Age, sex, and contraction protocol all affect the metabolic response to muscle contraction. We have previously examined fatigue and endurance in aged mice including restoring redox status associated with improved mitochondrial function in female mice [8, 9]. We have also shown that high-intensity stimulation can activate nuclear erythroid 2-related factor 2 (Nrf2) in both the stimulated and contralateral control leg [24]. These results led us to hypothesize that metabolic response to muscle contraction would be dependent on the type of protocol used. The two contraction protocols used here produced very different results in terms of metabolomic changes and oxidation of mitochondrial substrates between both female and males, but also revealed differences in response to contraction with age. Previous studies in both rodents [25, 26] and humans [27] have shown sex differences in myosin expression. Sexual dimorphism and differential myosin heavy chain expression has been implicated in divergent response to aging between male and female muscles [28]. In addition, sex-based difference in contractility and muscle kinetics exists between males and females in both rodents [29] and humans [30].

Results here are consistent with previous findings that males produced greater maximum force than females whereas female mice were more fatigue resistant. Interestingly, only females showed a difference in changes to fatigue based on age following HII, however this was largely driven by greater force decline with age in females than males. This is consistent with previous results showing that sex differences in skeletal muscle fatigue in mice are partially linked to expression of estrogen receptor-beta (ERβ) [31] and the estrogen-ERβ pathway functions to control muscle growth and regeneration in female mice [32, 33]. In contrast to our previous reports [8, 9] treatment with ELAM had no effect on fatigue in aged mice. This may be because HII used in this study was designed as an extreme, but much shorter protocol designed to elicit maximum molecular response to contraction, while previous protocols were more focused on testing muscle endurance over a greater length of time. The lack of difference in maximum force and fatigue with ELAM treatment here is worth noting because it indicates that differences in response to contraction are not driven by changes in the relative challenge to the muscle following ELAM treatment.

Sex differences in fatiguability between males and females [34] may at least be partially due to differences in substrate utilization during and following exercise and is likely exacerbated with age [35]. At rest and during exercise men and women utilize substrates for energetic demand at different rates [36]. We have previously identified declines in ATPmax and Phosphate/Oxygen/O in aged mice [6, 8, 9] consistent with energy deficits with age in humans [5, 11]. Although experiments in this study were designed to acutely alter metabolism of the contracted muscle, we found that LISS and HII altered substrate utilization in the non-stimulated gastrocnemius of both young and aged animals. While most exerkines previously identified are released due to whole body exercise [37] the data here suggests that even mild acute muscle contraction of a small muscle group can release a myokine capable of acting systemically on other muscles and possibly other tissues to alter metabolism. Previous studies in mice have shown that distinct serum metabolomic profiles exist between exercised and rested mice including decreases in circulating amino acids [19]. Data presented here show that distinct metabolomic profiles exist as well between HII and LISS in muscle (Figures 5 and 7 and Table 1). None of the metabolites significantly altered by HII or LISS identified here have been previously identified following acute bouts of exercise in mice [19]. There are four explanations for why this may be the case. One, the previous study used male mice and we used female mice for metabolomics because females showed more robust substrate utilization changes to muscle contraction. Two, this study used acute muscle contractions as opposed to whole body exercise. Three, this study used targeted instead of untargeted metabolomics increasing our power to identify specific metabolites but limiting testing of the entire metabolite pool. Four, metabolomics presented here are a direct measure of metabolites within skeletal muscle whereas circulating metabolites represent the pool of metabolites actively secreted and taken up for metabolism by tissues. Data here suggests that changes in the circulating metabolites following exercise do not adequately represent the tissue response to exercise/contraction. Most of the pathways altered following HII and LISS in young mice are not activated by HII or LISS in aged animals. In fact, only three pathways were changed in the aged animals following LISS: citrate cycle, butanoate metabolism, and synthesis and degradation of ketone bodies. Given the central nature of the citric acid cycle in metabolism of carbohydrates, proteins, and fats it is not surprising that this pathway remains activated in age [38]. Butanoate metabolism pathway changes are largely driven by metabolism of short chain fatty acids most commonly produced by intestinal fermentation by the gut microbiota [39, 40]; however, many molecules in this pathway are ultimately used in the production of ketone bodies [41]. Finally, ketone bodies contribute strongly to metabolism both during and after exercise as available carbohydrate sources are exhausted [42, 43]. The synthesis and degradation of ketone bodies pathway was changed in every comparison, apart from young LISS. This may be because LISS in young animals is not strenuous enough to exhaust available energy sources resulting in mobilization of ketone bodies. Despite increased stress and fatiguing contractions HII still only managed to significantly change five metabolic pathways in aged muscle. The metabolic pathways shown to be changed here by both HII and LISS in aged animals suggests an overall inability to respond to metabolic demands of muscle contraction with age.

**Table 1.**
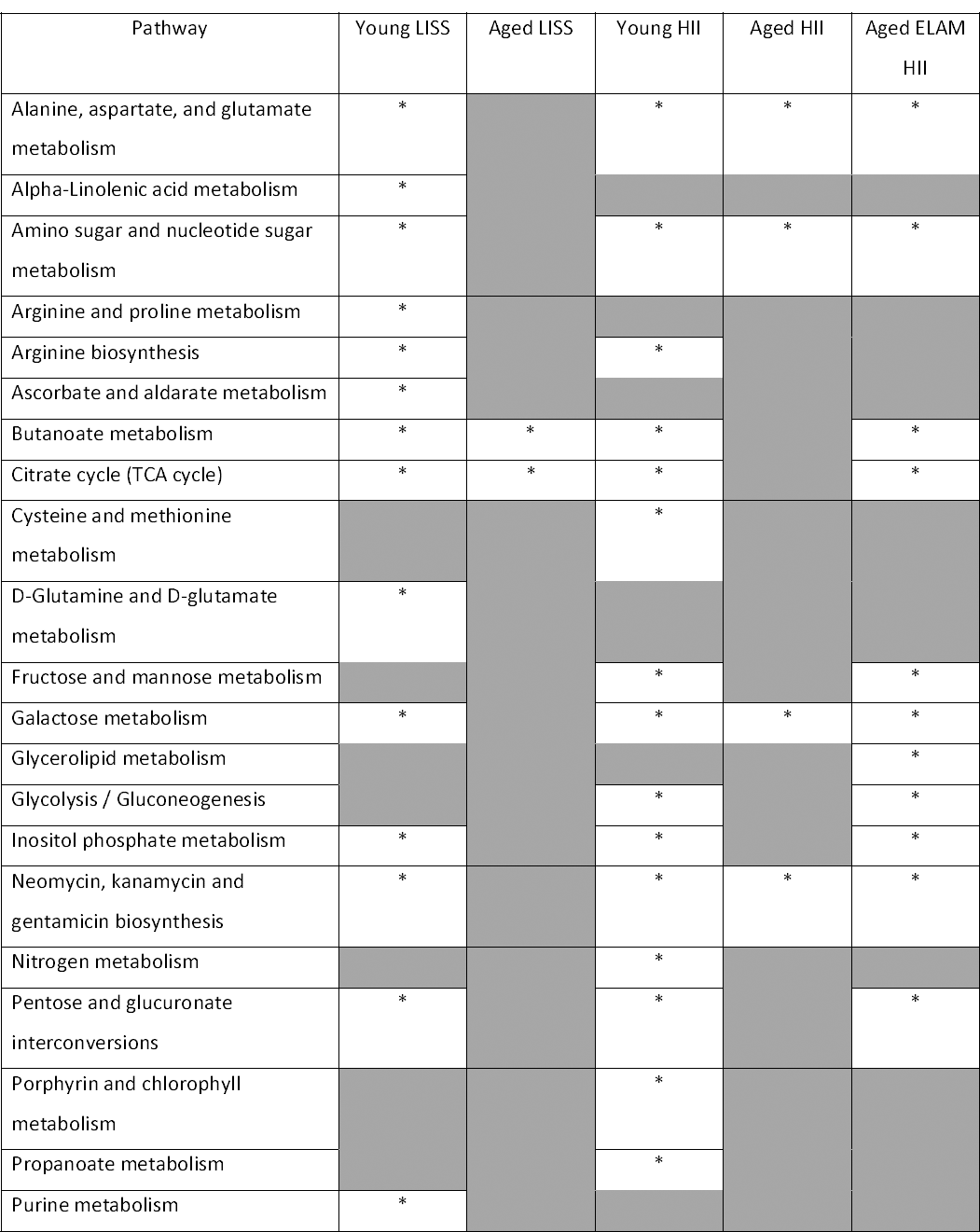
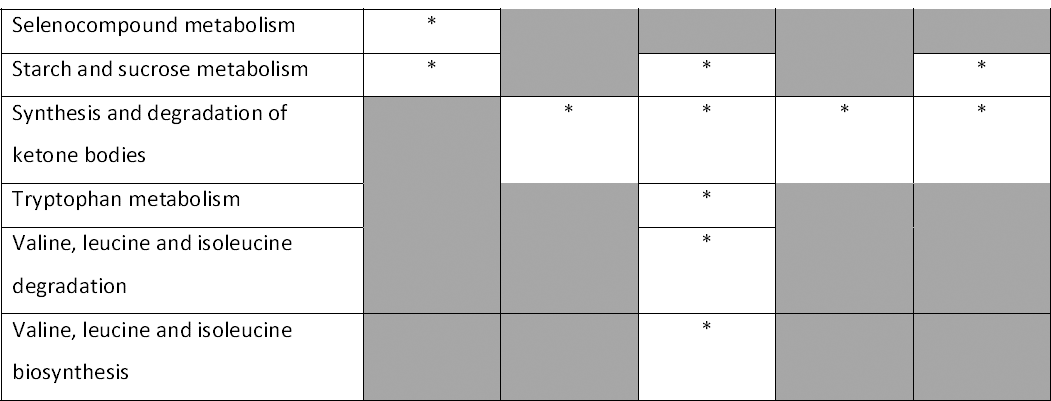
Metabolic pathway changes following HII and LISS. All pathways listed are significantly altered in at least one comparison. n=6-8, *-p<0.05

The reduced changes in specific metabolic pathways following both LISS and HII in aged comparisons may at least partially be due to deficient substrate utilization by the mitochondria in skeletal muscle following exercise. A large number of sugar and amino acid metabolism pathways are activated by contraction in young animals that are not present in the aged comparisons (Table 1). Intriguingly, inositol phosphate metabolism was present in both young LISS and HII comparisons and absent in aged LISS and HII; and was restored by treatment with ELAM. Inositol phosphate signaling has been shown to be a central component of energy maintenance [44] contributing to multiple signaling pathways and nutrient uptake processes including following muscle contraction [45]. The loss of activation of inositol phosphate metabolism with age following both stimulation protocols may be a contributing mechanism for poor metabolic response to muscle contraction. Deficient substrate utilization with age is further supported in aged female mice by a complete inhibition of oxidation of glutamate following HII. While circulating glutamate is normally associated with use for neurotransmission [46] skeletal muscle glutamate levels are altered in a number of pathological states [47]. Additionally, increased glutamate serum levels in sarcopenic patients positively correlates to greater function and muscle mass [48–50] suggesting glutamate has a central role in maintenance and metabolism of skeletal muscle with age. However, data here shows that glutamate metabolism can be inhibited following some bouts of intense exercise in females. This is particularly striking as glutamate has been shown as the only amino acid absorbed by skeletal muscle during exercise [51, 52]. Glutamate is a contributor to the citric acid cycle through its conversion to alpha-ketoglutarate by glutamate dehydrogenase (GDH), however, glutamate is also a necessary precursor for synthesis of the antioxidant glutathione [53]. In situations of increased oxidative stress such as following intense muscle contraction [54] or in the context of aging [9] it is likely that glutamate is preferentially used for glutathione synthesis rather than as a substrate for ATP generation. This does not adequately explain why we only noted inhibition of glutamate oxidation in aged females following HII contraction. In humans, serum levels of glutamate increase with age in females but not males [55] suggesting either an increased demand for circulating glutamate, or an inability to adequately uptake and/or metabolize circulating glutamate in females. Unfortunately, we were unable to directly measure GDH activity levels. This may have been because tissues were previously frozen inhibiting enzymatic activity, or it may have been due to lower levels of GDH in skeletal muscle relative to other tissues [56]. For future studies examining glutamate metabolism following exercise it would be informative to identify the contribution of glutamate to metabolism versus glutathione synthesis. Interestingly, glutamate inhibition following HII contraction extended even to the non-stimulated contralateral control leg suggesting some form of systemic exerkine signaling. Previous work in our lab has shown that high intensity *in vivo* contractions systemically activate Nrf2-mediated redox stress response [24]. Results here demonstrate that exerkine signaling further extends to control of mitochondrial function as well.

We were able to restore glutamate oxidation in aged females following HII by treating animals for 8 weeks with ELAM. We have previously shown ELAM improves *in vivo* bioenergetics [8, 9], increases endurance and fatigue resistance [9, 57], restores post-translational modifications [9, 58, 59], and rescues redox status in aged muscle [9, 60]. There are two potential mechanisms for ELAM restoration of glutamate oxidation. The first mechanism is through direct interaction of ELAM with glutamate dehydrogenase and/or glutamate metabolizing proteins to enhance glutamate utilization. ELAM has been shown to directly interact with four proteins involved in production of alpha-ketoglutarate [23]. While previously identified ELAM interactions have not been directly linked to increased glutamate metabolism this hypothesis is supported by decreased protein interactions with GDH in aged muscle that is linked to functional decline [61]. The second mechanism is through restoration of redox status by ELAM [9]. In aged muscle with decreased redox stress less glutamate would be needed for glutathione synthesis. This mechanism assumes a direct inhibition of glutamate oxidation with increased redox stress. However, data here supports this hypothesis and is consistent with our previous work showing ELAM increases reduced glutathione/oxidized glutathione both acutely [8] and long-term [9]. Additionally, GDH cysteine residue 376 has previously been shown to have increased protein S-glutathionylation with age that is reversed with ELAM treatment [9]. Work using molecular dynamics of the Bos taurus GDH analog has shown this residue resides in a key region of control for changes between the open and closed states of GDH during allosteric regulation of enzyme function [62]. Future studies focusing on whether redox sensitive post-translational modification of this specific cysteine residue exerts functional control over GDH may provide insight as to why HII can completely inhibit glutamate oxidation post-muscle contraction [8, 9]. The enhancement of glutamate oxidation following HII in ELAM treated muscle is supported by metabolomic analysis showing restoration of seven metabolic pathways activated in young muscle by HII that are lost in aged muscle (Table 1). Most of these pathways are related to amino acid and sugar metabolism suggesting that ELAM treatment increases muscle’s ability to respond to metabolic demand. Unfortunately, tests of additional substrates were not possible due to limited O2K chambers, but the data here strongly suggests that many metabolites are utilized in different ways based on age, sex, and even exercise protocol. Future studies examining metabolic flux should focus on the mechanisms of sexual dimorphism and changes with age.

## Conclusions

This study showed that female and male muscle have distinct metabolic responses to exercise that are also modulated by intensity of muscle contraction. The sex-dependent response to contraction extends into aged muscle as well. Metabolomics revealed that aged muscle is limited in its activation of metabolic pathways in response to the increased demand of contraction relative to young muscles. Further, improvement of mitochondrial function and redox status by ELAM can rescue glutamate utilization and restore metabolic pathways altered by HII following muscle contraction. This study is the first to identify inhibition of mitochondrial glutamate oxidation following muscle contraction and suggests active selection of glutamate for non-anaplerotic use following intense exercise in aged muscle.

## Methods

### Animals

All experiments in this study were reviewed and approved by the University of Washington Institutional Animal Care and Use Committee (IACUC). Female and male C57BL/6 mice were procured from the National Institute on Aging aged rodent colony. Young animals were between 5-7 months and aged animals were between 27-29 months old at the times of sacrifice. All animals were maintained on a 14/10 light/dark cycle at 21°C and given access to food and water *ad libitum* with no changes prior to experimentation.

### In vivo muscle contraction and mechanics

Animals were given O_2_ at 1 L/min and induced for anesthesia using 4% isoflurane. Once surgical plane of anesthesia was reached animals were moved to a water heated circulating platform maintained at 37°C, the right hindlimb was fixed in place at the knee and the foot was secured to a servomotor (Aurora Scientific, Aurora, ON, CA). During all procedures animals were maintained between 2-2.5% isoflurane. The gastrocnemius was stimulated via the tibial nerve using a high-power, bi-phase stimulator (Aurora Scientific) between 3-5 volts optimized for maximum force generation. Animals were stimulated with either HII (150 Hertz (Hz) every 3 seconds for six stimuli, followed by 10 seconds of rest, repeated for 10 bouts) or LISS (30 Hz every 10 seconds for 20 minutes). All data was analyzed using Dynamic Muscle Analysis Software (v 5.300 Aurora Scientific) and Prism 9.51. Maximum force comparisons were made using one-way ANOVA and a Tukey’s multiple comparisons test. Fatigue curves were compared using two-way repeated measures ANOVA with Šídák’s multiple comparisons test.

### Tissue dissection and partitioning

Immediately following *in vivo* muscle stimulation animals were euthanized using cervical dislocation. The stimulated and non-stimulated gastrocnemius were dissected and placed on ice. Each gastrocnemius was split into three parts. An approximate 3-6 mg portion of the red gastrocnemius was taken for mitochondrial respiration and the remaining muscle was uniformly split in two and snap frozen in liquid N_2_ for metabolomics or biochemical follow up assays.

### Mitochondrial respiration

Following dissection 3-6 mg of red gastrocnemius was separated into two fiber bundles and manually teased apart on ice in BIOPS (10 mM Ca-EGTA buffer, 0.1 µM free calcium,20 mM imidazole, 20 mM taurine, 50 mM K-MES, 6.56 mM MgCl_2_, 5.77 mM ATP, 15 mM phosphocreatine, pH 7.1) for 5 minutes or until visible fibers were loosely separated from adjacent fibers. Following manual teasing fiber bundles were permeabilized on ice in BIOPS with saponin (50 ug/ml) for 40 minutes with gentle rocking. Following permeabilization fibers bundles were washed for 5 minutes in BIOPS, followed by 5 minutes and 15 minutes in respiration buffer (RB, 0.5 mM EGTA, 20 mM taurine, 3 mM MgCl2, 110 mM Sucrose, 60 mM K-MES, 20 mM Hepes, 10 mM KH2PO4, 1mg/ml BSA, pH 7.1) on ice with gentle rocking. Following wash steps, fiber bundles were placed in RB in an Oxygraph 2-K dual respirometer/fluorometer (Oroboros Instruments, Innsbruck, AT) at 37°C, with 750 rpm stirring. Oxygen concentration was maintained between 250-450 uM. Respiration was stimulated with titrations to final concentration of 0.1 mM malate, 50uM ADP, 2.5 mM ADP, and 1 mM steps up to 10 mM glutamate; or with titrations to final concentration of 0.1 mM malate, 50 uM ADP, 2.5 mM ADP, and 1, 2, 3, 4, 5, 10, 20, 30, 40, 50, 60, and 70 uM palmitoyl carnitine. All data was analyzed using Datlab 7.4.0.4 (Oroboros Instruments) and Graphpad Prism 9.51. Respirometry values were compared using repeated measures one-way ANOVA with a Tukey’s multiple comparisons test.

### Metabolomics

Aqueous metabolites for targeted LC-MS profiling of 70 skeletal muscle samples were extracted using a protein precipitation method as previously described [63–65]. Samples were first homogenized in 200 µL purified deionized water at 4 °C, and then 800 µL of cold methanol containing 124 µM 6C13-glucose and 25.9 µM 2C13-glutamate was added (reference internal standards were added to the samples in order to monitor sample prep). Afterwards samples were vortexed, stored for 30 minutes at −20 °C, sonicated in an ice bath for 10 minutes, centrifuged for 15 min at 14,000 rpm and 4 °C, and then 600 µL of supernatant was collected from each sample (precipitated protein pallets were saved for BCA assay). Lastly, recovered supernatants were dried on a SpeedVac and reconstituted in 0.5 mL of LC-matching solvent containing 17.8 µM 2C13-tyrosine and 39.2 3C13-lactate (reference internal standards were added to the reconstituting solvent in order to monitor LC-MS performance). Samples were transferred into LC vials and placed into a temperature controlled autosampler for LC-MS analysis.

Targeted LC-MS metabolite analysis was performed on a duplex-LC-MS system composed of two Shimadzu UPLC pumps, CTC Analytics PAL HTC-xt temperature-controlled auto-sampler and AB Sciex 6500+ Triple Quadrupole MS equipped with ESI ionization source. UPLC pumps were connected to the auto-sampler in parallel and were able to perform two chromatography separations independently from each other. Each sample was injected twice on two identical analytical columns (Waters Xbridge BEH Amide XP) performing separations in hydrophilic interaction liquid chromatography (HILIC) mode. While one column was performing separation and MS data acquisition in ESI+ ionization mode, the other column was getting equilibrated for sample injection, chromatography separation and MS data acquisition in ESI-mode. Each chromatography separation was 18 minutes (total analysis time per sample was 36 minutes). MS data acquisition was performed in multiple-reaction-monitoring (MRM) mode. LC-MS system was controlled using AB Sciex Analyst 1.6.3 software. Measured MS peaks were integrated using AB Sciex MultiQuant 3.0.3 software. The LC-MS assay was targeting 361 metabolites (plus four spiked reference internal standards). Up to 182 metabolites (plus four spiked standards) were measured across the study set, and over 95% of measured metabolites were measured across all the samples. In addition to the study samples, two sets of quality control (QC) samples were used to monitor the assay performance as well as data reproducibility. One QC [QC(I)] was a pooled human serum sample used to monitor system performance and the other QC [QC(S)] was pooled study samples and this QC was used to monitor data reproducibility. Each QC sample was injected per every 10 study samples. The data were well reproducible with a median CV of 5.4 % over 2.5 days of non-stop data acquisition. Targeted metabolomics was examined using MetaboAnalyst 5.0 One Factor Statistical and Pathway Analysis. Features with >50% missing values were removed, and remaining missing values were excluded. Data was normalized using sample protein concentrations, mean-centered, and autoscaled. Metabolite changes were analyzed using paired students t-test between the contracted and non-stimulated muscle, and pathway changes were analyzed using enrichment analysis. All metabolite comparisons include a Holm-Bonferroni correction for multiple testing.

### Surgery and elamipretide treatment

Elamipretide treatment and surgical interventions were performed as previously described [9] delivering 3mg/kg/day of ELAM in osmotic minipumps (Alzet 1004) for 4 weeks followed by surgical removal and replacement using a fresh osmotic pump for an additional 4 weeks.

## Abbreviations

HII: high-intensity intervals
LISS: low-intensity steady state
GDH: glutamate dehydrogenase
ELAM: elamipretide

## Acknowledgements

We acknowledge The Northwest Metabolomics Research Center (NWMRC) at the University of Washington, Seattle, and NIH grant #1S10OD021562-01 that funded a purchase of the LC-MS platform used to acquire targeted metabolic profiling data. The authors would like to thank James MacDonald and Theo Bammler for their assistance with the organization and analysis of metabolomics.

## Supplemental Materials

**Supplemental Figure 1.**
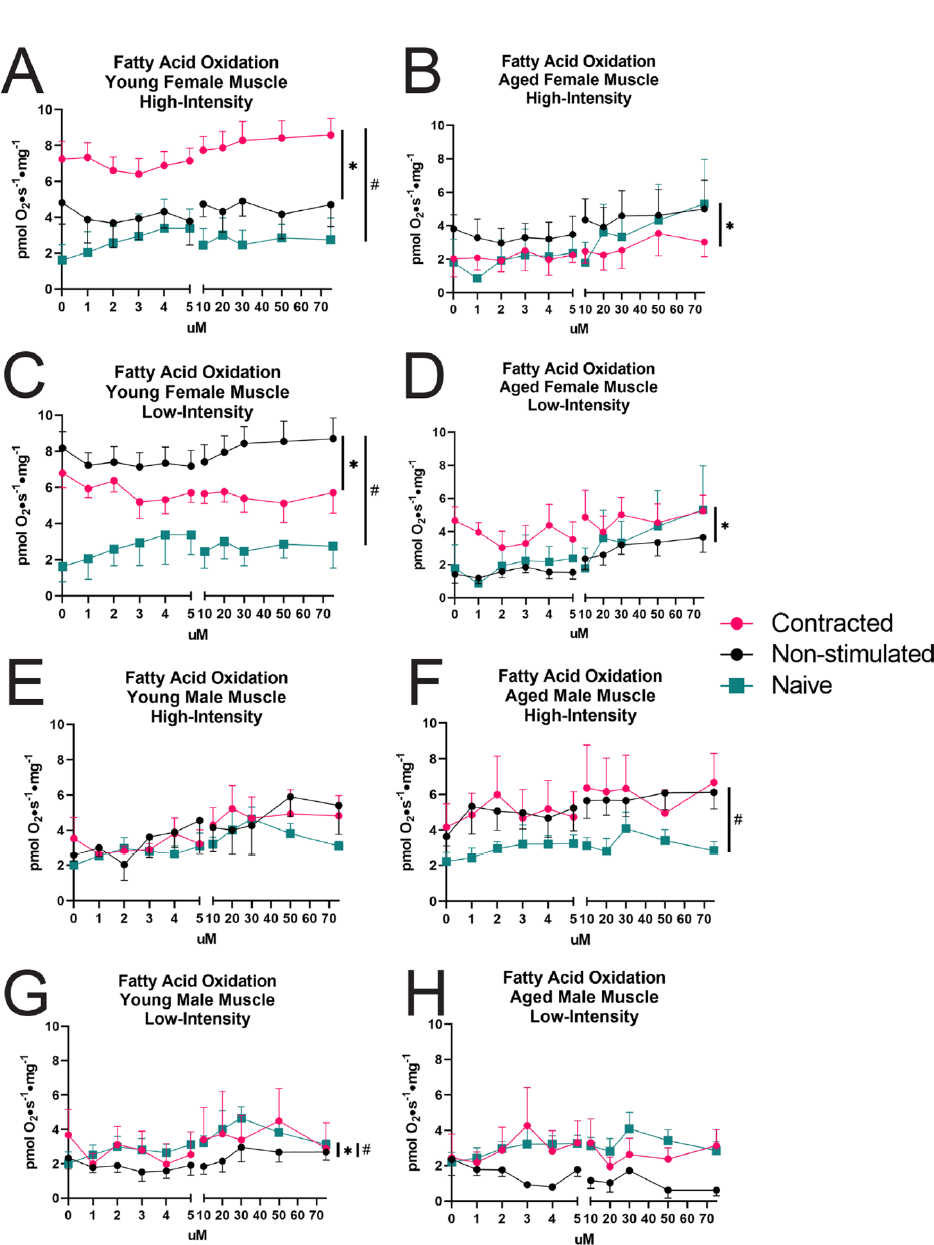
Palmitoyl carnitine utilization following acute muscle contraction. **A.** High-intensity intervals in young female mice, n=5 **B.** High-intensity intervals in aged female mice, n=7 **C.** Low-intensity steady state in young female mice, n=5 **D.** Low-intensity steady state in aged female mice, n=8 **E.** High-intensity intervals in young male mice, n=5 **F.** High-intensity intervals in aged male mice, n=9 **G.** Low-intensity steady state in young male mice, n=5 **H.** Low-intensity steady state in aged male mice, n=8 All data represented as mean ± S.E.M., *-p<0.05 repeated measures one-way ANOVA stimulated compared to non-stimulated, #-p<0.05 repeated measures one-way ANOVA effect compared to naïve.

